# Extrinsic repair of injured dendrites as a paradigm for regeneration by fusion

**DOI:** 10.1101/062372

**Authors:** Meital Oren-Suissa, Tamar Gattegno, Veronika Kravtsov, Benjamin Podbilewicz

## Abstract

Injury triggers regeneration of axons and dendrites. Research identified factors required for axonal regeneration outside the CNS, but little is known about regeneration triggered by dendrotomy. Here we study neuronal plasticity triggered by dendrotomy and determine the fate of complex PVD arbors following laser surgery of dendrites. We find that severed primary dendrites grow towards each other and reconnect via branch fusion. Simultaneously, terminal branches lose self-avoidance and grow towards each other, meeting and fusing at the tips via an AFF-1-mediated process. Ectopic branch growth is identified as a step in the regeneration process required for bypassing the lesion site. Failure of reconnection to the severed dendrites results in degeneration of the distal end of the neuron. We discover pruning of excess branches via EFF-1 that acts to recover the original wild-type arborization pattern in a cell-autonomous process. In contrast, AFF-1 activity during dendritic auto-fusion is derived from the lateral seam cells and not autonomously from the PVD neuron. We propose a model in which AFF-1-vesicles derived from the epidermal seam cells fuse neuronal dendrites from without. Thus, EFF-1 and AFF-1 fusion proteins emerge as new players in neuronal arborization and maintenance of arbor connectivity following injury in *C. elegans*. Our results demonstrate that there is a genetically determined multi-step pathway to repair broken dendrites in which EFF-1 and AFF-1 act on different steps of the pathway. Intrinsic EFF-1 is essential for dendritic pruning after injury and extrinsic AFF-1 mediates dendrite fusion to bypass injuries.

**Author summary:** Neurons in the central nervous system have very limited regenerative ability, they fail to remodel following amputation and only in some invertebrates, axons can repair themselves by fusion. Some genetic pathways have been identified for axonal regeneration but few studies exist on dendrite regeneration following injury. To determine how neurons regenerate dendrites following injury we study the *C. elegans* PVD polymodal neurons that display an arborized pattern of repetitive menorah-like structures. We injure dendrites by laser microsurgery, follow their fate and show that broken primary dendrites often regenerate via fusion. We describe how PVD dendrites regenerate and present roles for EFF-1 and AFF-1 proteins in fusion and remodeling of menorahs. Menorahs lose self-avoidance and AFF-1 fuses them, bypassing the injury site. Branch sprouting, EFF-1-mediated pruning, and arbor simplification completes regeneration. When auto-fusion fails the distal arbor degenerates. Surprisingly, AFF-1 acts non-cell autonomously to mediate dendrite fusion. We propose that extracellular vesicles derived from the lateral epidermis fuse severed dendrites in a process reminiscent of enveloped virus-mediated cell fusion without infection.

## Introduction

Sensory perception relies on networks of neurons that monitor and modify behavior to assure that animals are able to locate food, sense their environment and avoid predators or other threats [1]. This perception depends on the integrity and spatial-coverage of the receptive field [2]. Axonal and dendritic trees play an essential role in processing and transducing information to ultimately evoke the appropriate response of the organism. In the central nervous system (CNS) of adult mammals axon regeneration following injury is limited [3]. Therefore, the regenerative process following axon severing has been the focus of numerous studies [3–5]. It is believed that the main reasons why axons fail to regenerate are a reduction in neuronal growth capacity and inhibitory extrinsic factors. However, the molecular mechanisms of regeneration are not well understood. Recent studies have suggested that modulation of intrinsic neuronal activity by mammalian target of rapamycin (mTOR) and G-protein-coupled receptor (GPCR) signaling promote axon regeneration [4, 6]. In parallel there is evidence for a molecular pathway for axonal degeneration that affects regeneration [7]. The molecular mechanisms required for regeneration by regrowth following axonal injury are actively studied and numerous pathways have been identified [3–5, 8–12]. In contrast to regeneration by regrowth, a different strategy for axonal regeneration that has been observed in diverse invertebrates is reconnection by fusion of severed axons [13–18].

The nematode *C. elegans* is a powerful model to study neuronal regeneration after injury [19]. It has been recently found that injured axons of motor-and mechanosensory-neurons regrow and in some cases fuse after *in vivo* severing using laser surgery [15, 16, 18–21]. Moreover, screens for genes with roles in axon regrowth identified many genes required for axon regeneration [15, 22, 23].

Compared to axonal regeneration and degeneration pathways, much less is known about dendritic regeneration following injury [24–28]. Recent studies have identified the PVD and FLP neurons as highly branched bilateral neurons in *C. elegans,* which display a stereotypic arborization pattern composed of repetitive structural units known as menorahs (**Figures 1A and 1B**)[29–37]. The PVD is highly polarized with a single axon ventral to the cell body and complex but stereotyped dendritic arbors [33, 37] making it an ideal system to study different aspects of the generation, maintenance, regeneration and degeneration of dendritic trees. The PVD neurons are two polymodal nociceptors, responsible for an avoidance response generated after harsh mechanical stimuli to the main body or exposure to cold temperatures [1, 38, 39]. Animals in which PVD neurons are laser-ablated fail to respond to harsh touch [38]. Recent studies uncovered the degenerin ion channels DEG/ENaC, MEC-10 and DEGT-1 that sense harsh-touch, and the TRPA-1 channels that respond to cold temperatures [35, 40]. Moreover, researchers have identified numerous genetic pathways involved in dendritic arborization and maintenance of the PVD structure [33, 41–47].

**Fig 1.**
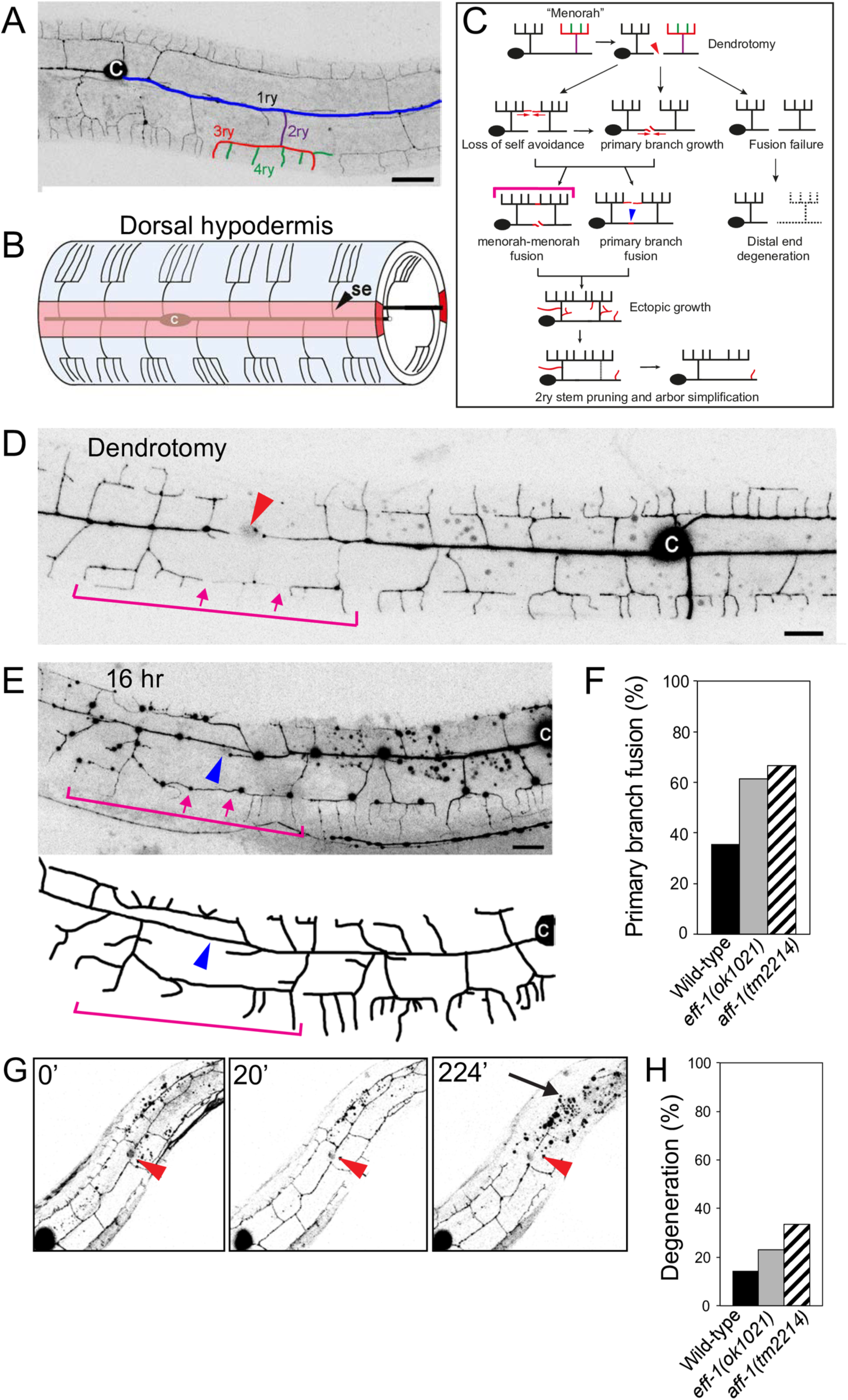
Dendrite regeneration of multibranched PVD neurons following laser microsurgery in *C. elegans*. **(A)** A wild-type animal expressing *DES-2::GFP*, illustrating the PVD neuron elaborate branching pattern (inverted image). Branches of one menorah are numbered primary to quaternary (1ry to 4ry), and color-coded: blue, purple, red, and green, respectively [33], c= cell body. **(B)** Schematic model of hypodermal cells and PVD menorahs in a young adult, left view. The wild-type PVDs grow between the hypodermis (outer cylinder, light blue) and the basement membrane of the hypodermis (not shown), extending processes that branch out to form the menorah structures. In light red is the left hypodermal seam syncytium. Modified from [33], se= seam cells, c= cell body. **(C)** Cartoon summarizing the different stages in PVD regeneration following injury. Two-photon dendrotomy (see Materials and methods) of the primary process (red arrowhead) leads to dynamic changes in the PVD arbor: loss of branch self-avoidance and growth is followed by primary branch fusion (blue arrowhead), menorah-menorah fusion (pink bracket), or both. There is an additional phase of dynamic growth and pruning, leading to arbor refinement. When the branches fail to fuse, the distal end undergoes degeneration. **(D-E)** Primary and menorah-menorah fusion following dendrotomy. L4 animal just after surgery (**D**) and 16 hr post-surgery (**E**). Red arrowhead marks the site of dendrotomy. Blue arrowhead marks reconnection site of dendrotomy. **(E)** The severed distal and proximal ends of the primary branch reconnected (blue arrowhead). Distal and proximal menorahs fused to form an additional connection between the soma and distal end (Pink arrows and bracket). **(F)** Percentage of primary branch fusion during regeneration in wild type (n=14), *eff-1(ok1021)* (n=13) and *aff-1(tm2214)* (n=12) dendrotomized animals. Differences are not statistically significant (Fischer’s Exact test). **(G)** Live imaging of an injured young-adult animal in which reconnection was unsuccessful, and degeneration of the distal arbor occurred. Time after injury is indicated in minutes. Red arrowhead– site of injury. Arrow– degenerating arbor. Images are taken from **Movie 3**. Posterior is up and dorsal is left. Scale bars represent 20 µm in (A) and 10 µm in (D-E). **(H)** Percentage of animals with degenerating distal end following dendrotomy in wild type (n=14), *eff-1(ok1021)* (n=13) and *aff-1(tm2214)* (n=12) animals. Differences are not statistically significant (Fischer’s Exact test).

The dynamic pathway of PVD arborization revealed an unexpected function of EFF-1 fusogenic protein in sculpting neuronal trees [33]. EFF-1 mediates epithelial and muscle cell-to-cell fusion [48–53], auto cell fusion in the digestive tract [54], axonal fusion following injury [15, 17, 18, 21] and acts in the PVD neurons to trim menorahs [33]. *eff-1* mutants have hyperbranched disorganized menorahs and EFF-1 cell-autonomous expression is sufficient to reduce the number of branches and to rescue disorganized menorahs [33]. EFF-1 controls dendritic plasticity via retraction of excess branches, by fusing branches and by forming loops that restrict further growth [33]. AFF-1, a paralog of EFF-1, mediates fusion of the anchor cell to form the utse/hymen, fuses the lateral epidermal seam cells and merges some embryonic epithelial cells [55]. In addition, AFF-1 is induced by Notch to auto-fuse a myoepithelial toroid [54], and fuses the excretory duct cell to form a single-cell tube [56, 57]. Here we determine a cellular pathway for dendritic remodeling following injury. We uncover the functions of two fusion proteins, EFF-1 and AFF-1, in different stages of the regeneration of dendritic arbors of the PVD polymodal neuron in *C. elegans*.

## Results

### Dissection of dendritic regeneration in *C. elegans* mechanosensory neurons

The regenerative ability of axons following injury has been previously described in vertebrates and invertebrates [14, 19, 58–61]. The morphological and molecular changes that occur following dendritic amputation remain mostly unexplored [24–27, 33, 62, 63]. To study the process of regeneration of the PVD dendrites following amputation we performed dendrotomy of arborized neurites using a femtosecond laser [8–15, 16, 20]. A successful regeneration was defined as a process in which the severed branch was able to reconnect with its target [64]. Failure to rejoin the two parts of the severed primary dendrite results in degeneration of the distal part and in some cases a complete degeneration without regrowth of the multi-menorah dendritic tree. Temporal analysis of PVD dendrite dynamics following injury revealed several overlapping steps in arbor regeneration (**Fig 1C**). PVD dendrites showed robust regeneration when severed at the L4 stage. We found that severed primary dendrites grew towards each other (**Movies 1 and 2)** and in around 40% of the animals were able to reconnect the distal end to the soma via fusion (**Fig 1D and 1E**).

It was shown that following axotomy of the PLM mechanosensory neuron, reconnection of the axon to the distal branch is dependent upon EFF-1, but not AFF-1 activity [17, 18, 21]. In contrast, we found that primary dendrotomies in *eff-1* or *aff-1* single null mutants were repaired and regeneration via primary branch fusion occurred (**Fig 1F**). Due to the sub-viability of *eff-1aff-1* double mutants [55], we were unable to test whether there is redundancy between these genes in primary branch fusion. It is conceivable that there is redundancy in the fusion machinery that fixes broken neurites or that an unidentified fusogen is required for postembryonic primary dendrite auto-fusion after microsurgery. Failure to reconnect following dendrotomy resulted in degeneration of the distal end and took place within 12 hr in about 20% of the operated animals (**Fig 1G**, arrow, **1H** and **Movie 3**). Thus, following primary branch microsurgery the two ends grow towards each other, the tips meet, connect and the integrity of the distal arbors is maintained.

### Dendrotomy reveals menorah plasticity and causes loss of self-avoidance

The dendritic architecture of the PVDs is maintained by a contact-dependent self-avoidance mechanism. The tertiary branches withdraw upon contact of a neighboring branch, maintaining the menorah architecture [34, 43]. To test whether self-avoidance is maintained after injury, we explored the spatial dynamics of regenerating dendrites. Two hours after injury, we observed tertiary branches from neighboring menorahs that contacted each other and extended far from their initial location, resulting in a structure of overlapping menorahs (**Fig 1D and 1E**, bracket). Some of these overlaps extended and occurred between menorahs originating from both sides of the lesion (**S1 Fig, brackets and Movies 1 and 2**). These overlapping structures persisted even 48 hr after the injury (**S1F Fig**). Dendrotomy at earlier stages, such as the L3 stage, showed similar results; animals exhibited loss of avoidance-mechanisms and branch overlap (data not shown). These results suggest that upon injury the avoidance mechanisms are lost, making it more likely that a new connection will form to compensate for the injury.

### Menorah-menorah fusion bypasses the severed primary dendrites

We have previously shown that during wild type development PVD and FLP terminal quaternary branches can auto-fuse with one another to maintain menorah structure and to limit further growth [33]. To determine whether fusion of terminal dendrites is part of the regeneration process, we analyzed the overlapping branches after injury and found that most of the reconnections bypassed the injury site through menorah-menorah fusion, resulting in giant menorahs (**Fig 2A-2D, and S2B Fig**, brackets). To judge the connectivity of the tertiary branches we verified that the distal processes do not degenerate, and analyzed GFP-signal continuity using confocal microscopy and live imaging (**Fig 2D**, **Movies 1, 2 and 4**). In addition we used a photoconvertible Kaede cytoplasmic reporter expressed in the PVD [65] to demonstrate that the menorahs have fused and therefore have connected to bypass the lesion site (**S3 Fig**). We found that in animals where menorah-menorah fusion took place the distal fragment did not degenerate, regardless of primary-primary branch reconnection (n=23). Thus, terminal branch auto-fusion acts as a mechanism to bridge the gap between the PVD soma and the distal end to maintain connectivity and avoid degeneration.

**Fig 2.**
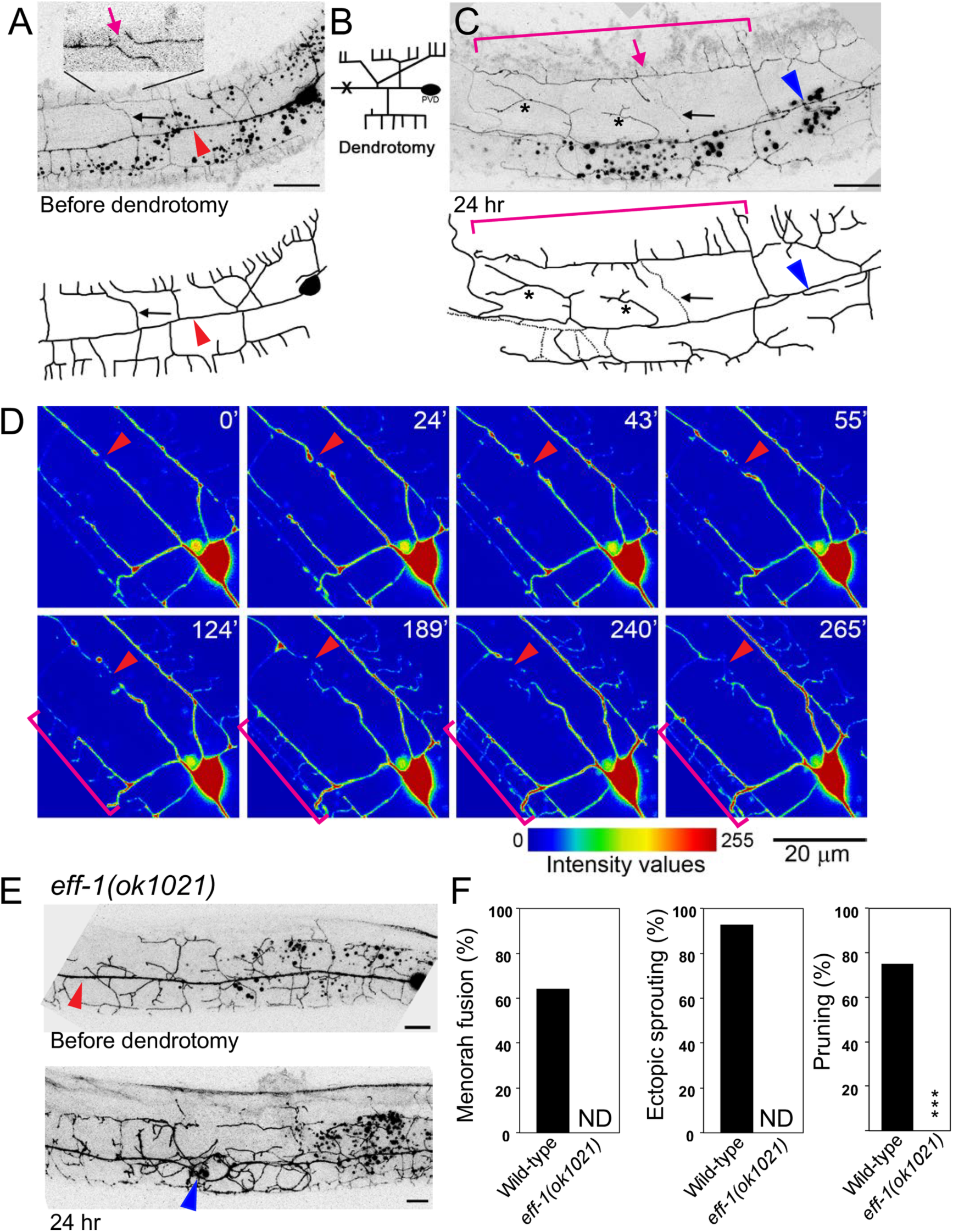
The dynamic pathway of PVD dendrite regeneration after injury. **(A-C)** Dendrotomy of an L4 animal resulted in menorah-menorah fusion. **(A)** Just prior to the injury separate menorahs can be seen (pink arrow in inset). Red arrowhead points to the lesion site. **(B)** Schematic showing the site of injury (marked with an X). **(C)** Formation of giant menorahs (pink bracket) and ectopic sprouting **(asterisks)** within 24 hr. Pink arrow points to the site of loss of self-avoidance. Blue arrowhead points to the site of 1ry-1ry fusion. Pruning of secondary branches also occurs (black arrow, dotted lines, also marked in (**A**)). **(D)** Intensity values view of images from a time-lapse movie of injured L4 wild-type worm (**Movie 4**). Time after injury is shown at the upper right corner in minutes. Arrowheads point to injury site and pruning of branches. Brackets mark two menorahs from distal and proximal ends bypassing the break and contacting one another. **(E)** *eff-1* null animal before and 24 hr after dendrotomy. There is a successful reconnection of the severed primary branch (arrowhead) and pruning failure (the arbor does not undergo simplification). Scale bars represent 10 µm. **(F)** Dendrotomy-induced phenotypes in *eff-1(ok1021)* animals. Menorah-menorah fusion, percentage of animals that contained fused giant menorahs; ectopic sprouting, fraction of animals showing growth of additional processes; pruning, percentage of animals in which PVD underwent branch refinement after fusion. In *eff-1(ok1021)* mutants there is excess branching and disorganized branch structure [33], thus we could not determine (ND) whether menorah-menorah connections and ectopic sprouting occurred. *\*\*\*P*<0.001, Fisher’s exact test. n values: 14, 13 animals for wild-type and *eff-1(ok1021)*, respectively.

### Sprouting, pruning, and arbor simplification complete regeneration

We observed that the PVD dendrites appeared highly dynamic after injury, showing ectopic growth of terminal branches. These results suggested that growth is not restricted to a subset of branches, and can occur throughout the neuron, allowing for massive regeneration (**Fig 2C and S1 Fig**; asterisks). To verify that growth is stimulated specifically due to the dendrite injury and not because of laser damage, we examined mock-injured animals for PVD morphology changes. We injured animals near the PVDs, at the same focal plane, but without hitting any dendrite. PVD mock-operated animals showed normal growth with no excess sprouting or changes in the PVD morphology (data not shown). These results demonstrate that dendrite severing specifically induced terminal branch reconnection by fusion, ectopic sprouting and regeneration (**Fig 1C**, **Movies 1 and 2**).

Interestingly, some of the distal secondary stems were eliminated following the reconnection, leaving just one or two secondary stems per giant menorah (**Fig 2C and S2B Fig**; arrows). These dendritic rearrangements persisted even 48 hr after surgery, leading to simplification of the dendritic trees and leaving mainly giant menorahs. Thus, active elimination of excess branches occurs in order to recreate a pattern resembling wild-type menorahs. Analysis of time-lapse movies showed that excess branches were eliminated, and pruning occurred concomitantly with growth around the injury site (**Fig 2D** and **Movie 4**). Taken together, the PVD dendrites are able to successfully regenerate following dendrotomy, inducing dynamic remodeling by branch growth and elimination (**Fig 1C**).

### EFF-1 is essential for pruning excess branches after dendrotomy

The central function of EFF-1 in PVD developmental arborization is in quality control trimming of excess and abnormal branches [33]. To determine whether EFF-1 acts in simplification following injury we amputated primary dendrites in *eff-1* mutants and followed the repair process. We found that *eff-1* mutants maintained hyperbranched and disorganized menorahs and failed to simplify the dendritic tree following injury (**Fig 2E**). These phenotypes suggest that *eff-1* acts in branch retraction and simplification induced by severing of the primary branch. We were not able to determine whether *eff-1* participates in menorah-menorah fusion because the hyperbranched and severely disorganized arbors prevented us from identifying menorah fusion and additional ectopic sprouting (**Fig 2F**). In contrast, both uncut and dendrotomized *eff-1* mutants showed no pruning, demonstrating that *eff-1* is required for branch simplification following dendrotomy. In addition, cell-autonomous expression of EFF-1 in the PVD resulted in excess pruning [33, 66]. Thus, EFF-1 acts cell autonomously to simplify excess sprouting following dendrotomy and is sufficient to trim branches and simplify arbors.

### AFF-1 is required to bypass cut dendrites via menorah-menorah fusion

Since the *C. elegans* known fusogens, EFF-1 and AFF-1, are essential and sufficient to fuse cells in *C. elegans* and heterologous cells in culture [50, 51, 55, 67, 68], we hypothesized that they may be required to regenerate broken neurites by homotypic fusion.

Because EFF-1 prunes dendrites by branch retraction [33, 66] we decided to determine a possible role for *aff-1* following dendrotomy, we asked whether menorah-menorah fusion occurs in *aff-1* injured-mutants. We found that while in *aff-1* mutants dendrite development was normal (**Fig 3A**), following dendrotomy most of the reconnections were between the re-growing primary dendrite and its distal fragment, rather than through menorah fusion as in wild-type animals (**Fig 3D**). This observation suggests a fusogenic function for AFF-1 in terminal branch fusion. In *aff-1* mutant animals we found some exceptions in which some menorahs overlapped, but we did not observe fusion between menorahs followed by secondary stem degeneration (**Fig 3D**). In dendrotomized *aff-1* animals failure to rejoin the dendritic trees resulted in degeneration of the distal part of the arbor (**Fig 3A and 3D**). Thus, while *aff-1* has no apparent role in normal PVD arborization, it is required for terminal branch fusion following dendrotomy.

**Fig 3.**
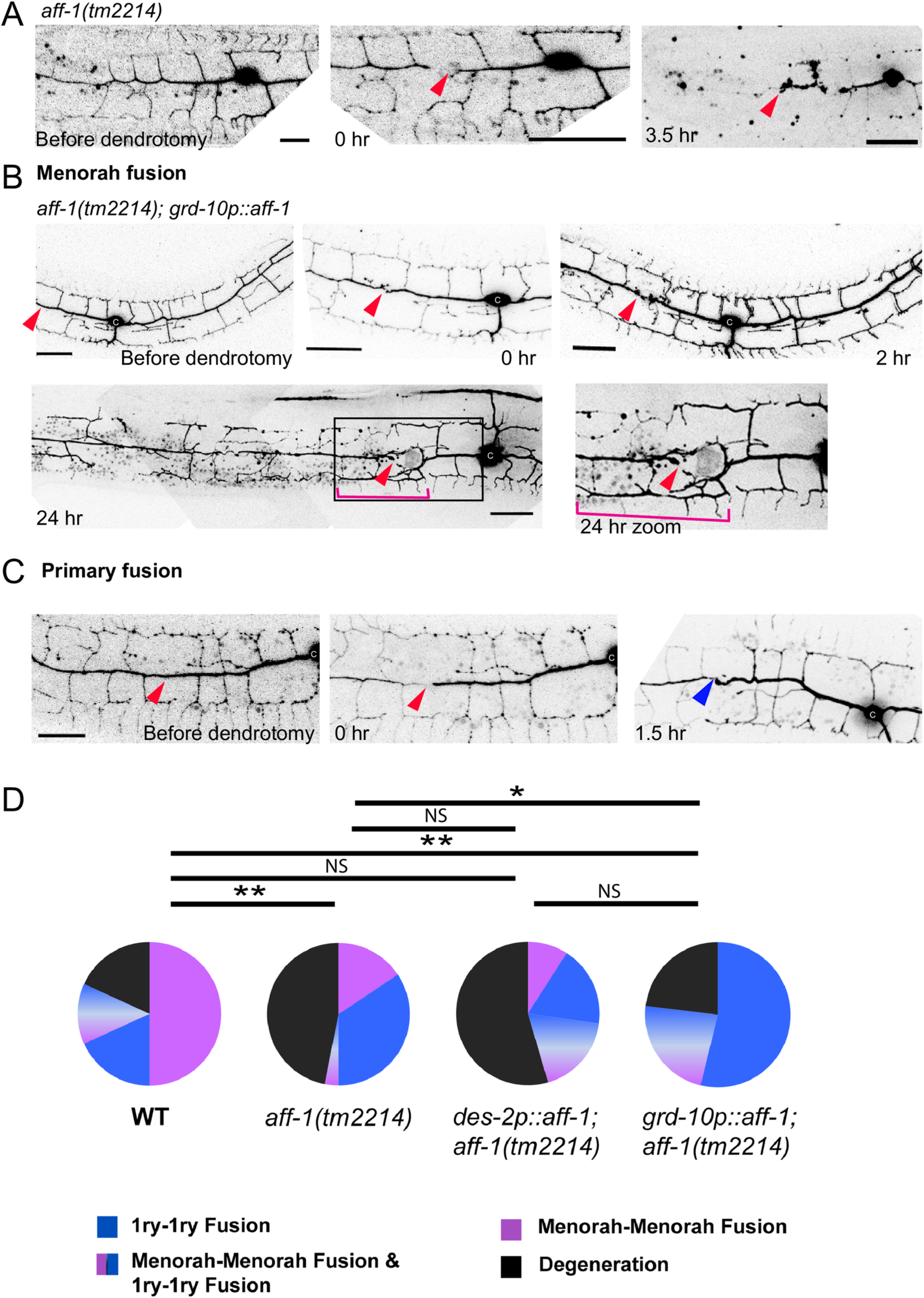
*aff-1* dendritic reconnection patterns are rescued cell non-autonomously. **(A)** Dendrite regeneration following PVD nanosurgery in *aff-1(tm2214)* mutant animal. PVD is shown before and after nanosurgery (t=0 and 3.5 hr). In non-injured *aff-1* mutant animals PVD branching pattern is unaffected. After injury, fusion does not occur and the distal processes undergo degeneration (t=3.5 hr). **(B)** Dendrite regeneration following PVD nanosurgery in *aff-1(tm2214); grd-10p::aff-1* animals. PVD is shown before and after nanosurgery (t=0, 2, and 24 hr). PVD reconnection occurred through fused menorahs (pink bracket). Red arrowhead marks site of injury. Primary branches did not fuse (red arrowhead). **(C)** Dendrite regeneration following PVD nanosurgery in *aff-1(tm2214); grd-10p::aff-1* animals. PVD is shown before and after nanosurgery (t=0 and 1.5 hr). PVD reconnection occurred through primary fusion (blue arrowhead). **(D)** PVD post-injury outcomes displayed in color coded pie graphs as Magenta– Menorah-Menorah fusion, Blue– primary-primary fusion, Magenta and Blue– Menorah-Menorah fusion and primary-primary fusion, Black– degeneration. Wild type n=22, *aff-1(tm2214)* n=32, *aff-1(tm2214);des-2p::aff*-n=11, *aff-1(tm2214); grd-10p::aff-1* n=13. ** *P*<0.01; * *P*<0.05; NS *P*>0.05. Dendrotomy site – red arrowhead, fused menorah – pink bracket, primary fusion – blue arrowhead. Scale bars represent 20 µm. Statistics was calculated using the Freeman-Halton extension of the Fisher exact probability test for a two-rows by four-columns contingency table, http://vassarstats.net/fisher2x4.html.

AFF-1-mediated membrane fusion of terminal branches emerges as the main mechanism by which dendritic repair occurs in *C. elegans*. We were not able to detect *aff-1* expression in the PVD neither before nor after dendrotomy. To determine whether AFF-1 acts extrinsically to the PVD to reconnect dendrites we attempted to rescue menorah-menorah fusion in *aff-1* mutant using expression of *grd-10p*::AFF-1 in the epithelial seam cells. We found induced recovery of reconnection and reduced degeneration to wild-type levels (**Fig 3B-3D**). In contrast, we found that PVD expressing *des-2p*::AFF-1 was not able to rescue the menorah-menorah fusion failure phenotype (**Fig 3D**). Expression of *dpy-7p::aff-1* from the hypodermis was toxic in *aff-1* homozygous mutants and did not significantly improved PVD regeneration in heterozygous *aff-1/+* animals, suggesting that non-autonomous expression only from epithelial seam cells is sufficient to improve the ability of the PVD to rejoin the branches by fusion (**S4 Fig**). The non-cell autonomous activity of AFF-1 in PVD regeneration by auto-fusion was unexpected since for cell-cell fusion AFF-1 acts homotypically in both fusing plasma membranes [68].

### AFF-1-containing extracellular vesicles may repair the PVD by fusing with it

To determine AFF-1 expression and localization before and after dendrotomy we imaged AFF-1 in worms expressing mCherry in the PVD, using a 30kb fosmid-based GFP reporter [69]. We could not detect AFF-1 in the PVD at any stage during development or following dendrotomy. Instead, AFF-1 is strongly expressed on the plasma membrane, filopodia and internal puncta in the lateral seam cells. This was expected since the seam cells fuse homotypically between the L4 and adult molt via AFF-1-mediated fusion [55]. Using structured illumination microscopy we found extracellular puncta containing AFF-1::GFP that apparently were derived from the seam cells (**Fig 4A**, arrowheads). Using live spinning disk confocal microscopy we found that the vesicles containing AFF-1::GFP were observed outside the seam cells in control animals that were not dendrotomized (**Fig 4B, Movies 5, 6 and 9**).

**Fig 4.**
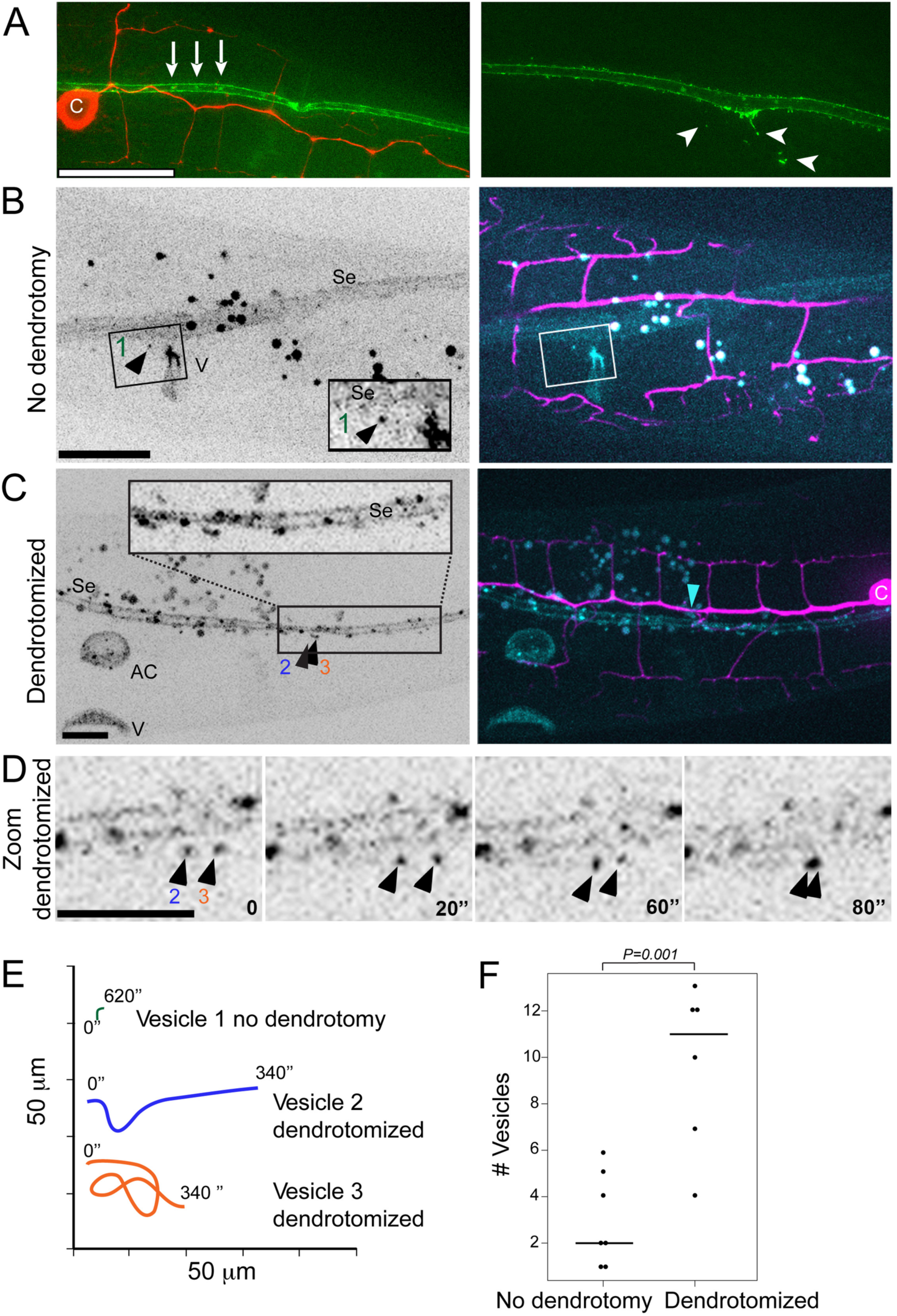
AFF-1 protein dynamics during PVD dendrite regeneration. **(A)** Superresolution structured illumination microscopy (SIM) image of aligned *AFF-1::GFP* (green) vesicles in the seam cells (left, white arrows) and *AFF-1::GFP* extracellular vesicles derived from the seam cells (right, white arrowheads) in intact wild type animals. c, cell body. PVD is labeled in red. **(B)** Spinning disk confocal images of *AFF-1::GFP* expression inside and outside the seam cells (Se), and in the vulva (V) of an intact L4 animal (left). *AFF-1::GFP* (cyan) expression in the seam cells and in vesicles near the seam cells is shown in a two channels merged image (right; PVD in magenta; **Movie 5**). Inset shows magnification of a single *AFF-1::GFP* vesicle near the seam cells (left, black arrowhead, numbered 1; **Movie 6**). **(C)** *AFF-1::GFP* expression inside and outside the seam cells (Se), in the anchor cell (AC) and in the vulva (V) of a reconnected PVD L4 animal (left) is shown 2 hr post-dendrotomy in a spinning disk confocal image. *AFF-1::GFP* is expressed in vesicles inside and outside of the seam cells, in the anchor cell (AC) and in VulA ring (see **Movie 7;** right, blue arrowhead, site of reconnection). Inset shows magnification of area with multiple vesicles, two are labeled (2,3) and shown in panels **(D-E)**. **(D)** Two *AFF-1::GFP* vesicles, numbered 2 and 3, are moving near the seam cells in a PVD-dendrotomized animal. Four time points are shown (0, 20, 60 and 80 sec; **Movie 8**). **(E)** Vesicles dynamics (1, 2 and 3) is demonstrated in a color-coded graph during 340 seconds for vesicles 2 and 3 and 620 seconds for vesicle number 1. The plot illustrates vast differences between *AFF-1::GFP* vesicles movements in animals with an intact PVD versus *AFF-1::GFP* vesicles in PVD dendrotomized animals (**Movies 6 and 8)**. **(F)** Dot Plot of the number of *AFF-1::GFP* vesicles outside *aff-1* expressing organs in animals with intact PVD versus in animals with dendrotomized PVD. The graph demonstrates statistical significant difference between the mean number of *aff-1::GFP* vesicles outside *aff-1* expressing organs such as seam cells, vulva, anchor cell and the uterus. *P*=0.001, statistics was calculated using the nonparametric Mann-Whitney test.

Following dendrotomy the AFF-1::GFP signal in the seam cells was brighter (**Fig 4C**) and there was a five-fold increase in the mobility and number of extracellular vesicles (**Fig 4D-4F, Movies 7, 8, 10 and 11**). Thus, taken together our results show that AFF-1 regenerates severed PVD dendrites in a surprisingly non-cell autonomous way from the seam cells.

## Discussion

### Hypothesis: AFF-1-extracellular vesicles merge injured neurons from without

Cell-cell fusion from within occurs when a fusogenic protein (e.g. a viral fusion protein following infection) or an endogenous cellular fusion protein (e.g. EFF-1) is expressed intrinsically in cellular compartments, including the plasma membrane. Fusion from without occurs when a viral particle fuses target cells without infecting them [70–72]. Here we discovered an example of fusion from without during neuronal regeneration. We propose that epidermal seam cells shed extracellular vesicles that travel 1000 nm or less to reach the PVD severed dendrites and the menorahs. These vesicles contain AFF-1 and can fuse dendrites from without (**S5 Fig**). AFF-1-containing vesicles derived from the lateral epidermal seam cells mediate fusion of severed dendrites and menorah auto-fusion to bypass the injury and to maintain dendritic tree structure and function. Extracellular vesicles (EVs, microvesicles, exosomes and ectosomes) from different sub-cellular and tissue origins have been proposed as vehicles for cell-cell communication during normal physiology, participate in the immune response, control coagulation and promote metastatic cancer [73]. These extracellular vesicles have been shown to exist in bacteria, archea, plants, fungi and animals [74–78]. In *C. elegans* EVs derived from ciliated neurons affect mating behavior and communication between animals [79]. Signaling EVs derived from the sperm activate oogenesis and ovarian muscle contraction [80]. EVs also participate in engulfment of dead cells [81] and morphogenesis of the embryo [82].

EVs are probably universal but diverge in size, shape and place of origin. They contain lipid bilayers, transmembrane proteins and nucleic acids. One of the characteristics of these EVs is their ability to fuse to target cells and deliver RNAs, plasmids, toxins and signaling molecules. However, the fusion proteins necessary to deliver and merge the diverse EVs have not been identified and characterized in any system. Mammalian cells transfected with *C. elegans* AFF-1 produce extracellular vesicles that have been biochemically and ultrastructurally characterized [68, 83]. Moreover, AFF-1-containing vesicles and pseudotyped particles are able to fuse to mammalian cells expressing EFF-1 or AFF-1. Thus, AFF-1 can mediate fusion of extracellular vesicles to cells expressing EFF-1 on the plasma membrane in a tissue culture system [68, 83]. Here we provide the initial evidence for a proposed mechanism that can fuse EVs to target neuronal cells in vivo. Surprisingly, these EVs can cause auto-fusion from without mediated by AFF-1 transmembrane fusion protein on their surface. Moreover, in our working model, these AFF-1-EVs derived from the *C. elegans* lateral epithelia can fuse neurons in vivo, thus directly promoting regeneration (**S5 Fig**).

### Neurodevelopmental genetic stages in dendrite repair after injury

The primary mechanism of PVD dendritic regeneration can be divided into five stages: (1) reattachment at site of injury, (2) loss of self-avoidance between adjacent menorahs, (3) menorah-menorah fusion to bypass lesions, (4) sprouting of compensatory branches and (5) pruning of excess branches (**Fig 1C**).

The interplay between two effector fusogens revealed a genetic pathway that links membrane remodeling during normal development and following neuronal injury. Here we focused on two cellular stages: Extrinsic stage (3) AFF-1-mediated menorah-menorah fusion from without and intrinsic stage (5) EFF-1-mediated trimming of excess branches from within. (**Figs 1C** and S**5**).

### AFF-1 merges terminal branches to bypass broken dendrites

Axonal fusion after injury is crucial for reestablishing synaptic contacts, to prevent degeneration, and for regaining neurological functionality [19, 64]. Although *eff-1* mutants failed to fuse broken axons [21], *eff-1* mutants succeed to merge injured dendrites. We tested the two known *C. elegans* fusogens, EFF-1 and AFF-1, as well as the EFF-1 paralog, C26D10.7 ([51, 84] and Oren-Suissa and Podbilewicz, unpublished results). We found that terminal-branch fusion following dendrotomy was significantly reduced in *aff-1* mutants compared to wild-type. However, none of these genes was independently required for primary dendrite fusion following an injury, suggesting either redundancy or that yet another *C. elegans* fusogen awaits identification. Based on these observations, we conclude that *aff-1* is required to heal dendritic wounds, specifically via menorah-menorah fusion.

Our data support a model in which *aff-1* and *eff-1* expression is highly regulated in the PVD. Following dendrotomy *eff-1* expression appears to be repressed allowing ectopic sprouting of terminal branches and loss of self-avoidance culminating with AFF-1-mediated menorah-menorah fusion via a surprising non-autonomous mechanism of fusion from without (**S5 Fig**).

### EFF-1 retracts and simplifies excessive branching following injury

The development of the nervous system includes the formation of many new branches, yet some of them are later eliminated. This process is termed pruning and is crucial for the formation of stable connections [85–90]. In the central and peripheral nervous systems, axons that are severed from their cell bodies degenerate by the genetically active process of Wallerian degeneration (WD) [7]. WldS mutant mice present slow WD and the persistent axon fragments appear to prevent regeneration by regrowth [7]. In addition, the ubiquitin/proteasome system acts to regulate axon degeneration in Drosophila, in vitro and in mice [91]. Thus, failure to remove axon debris following injury may physically block regeneration by regrowth of severed axons and WD appears to be required for efficient regeneration by regrowth [92]. Developmental pruning occurs in *C. elegans* AIM interneurons, and this process is regulated by the helix-turn-helix transcription factor MBR-1 [93] and Wnt-Ror kinase signaling [86]. Here, we demonstrated a novel mechanism of branch pruning following injury. We have previously shown that significant refinement is required to maintain the complex web of dendrites in the final state of perfectly sculpted PVD menorahs [33]. This dynamic nature persists following dendrotomy, and many new branches sprout around the injury site to ensure a successful connection will occur. This burst of growth is followed by the elimination of excess branches, and the arbor is simplified to resemble the original tree. This process is normally repressed probably by transcriptional mechanisms. Following injury and a stimulation of ectopic branching, *eff-1* is activated to simplify the arbors and regenerate a normal structure of menorahs. These results suggest that there is conservation of the molecular machinery for neurite refinement, and that the ability to alter existing morphologies is a key component in the development and survival of nervous systems.

### Is auto-fusion an alternative pathway to repair severed neurons?

Fusion of severed axons occurs in invertebrates for example in Aplysia [13], crayfish[14] and *C. elegans* [15–18] but rarely in vertebrates [6, 94]. Why is this the case and how can our study help us understand this? Invertebrates and vertebrates do have conserved pathways to regenerate injured branches via regrowth [3, 4, 9, 15, 95–97]. However, it appears that fusion of broken neurites or bypassing the injured site using fusion instead of rebuilding complex trees is a more energetically economical process. The use of extracellular vesicles could be a useful strategy to stimulate repair of injured branches in the CNS of vertebrates that usually cannot regenerate.

### Severed neurites can reconnect by suspending self-repulsion mechanisms

We have found that in *C. elegans* PVD, following dendrotomy there is a transient loss of self-avoidance between the menorahs (tertiary branches) that allows the reconnection by merging the menorahs and bypassing the site of injury, thus maintaining the dendritic trees. This is consistent with studies in leech embryos showing that laser microbeam severing of neurites of mechanosensory neurons result in that the detached branch stopped being avoided by the rest of the cell. This is consistent with a mechanism that controls self-avoidance and that requires physical continuity between the neurites [98]. In *C. elegans* this mechanism appears to involve netrins [43]. In contrast, in zebrafish detached fragments continue to repel the parent arbor [92]. Thus in zebrafish and probably in other vertebrates it is required to have a WD-like mechanism to remove fragments of sensory neurites before the process of regrowth can occur. It would be useful to find ways to induce merging of the severed neurites as occurs in some invertebrates.

Spinal cord injuries, experimental axotomies, surgical accidents, and diverse forms of neurodegeneration are all conditions that currently cannot be generally repaired [3, 6, 60, 61, 99]. Unveiling the mechanism of intrinsic *eff-1*-mediated dendritic simplification and extrinsic *aff-1*-mediated neuronal auto-fusion from without may pave the way for overcoming neurodegenerative diseases or injuries. The dynamic dendritic plasticity of the PVD neurons and the power of genetics in *C. elegans* along with its amenability for RNAi-based analysis and genome editing will uncover genetic pathways that will enable the study of additional mechanisms involved in PVD arborization during development and following injuries. In *C. elegans* AFF-1-containing vesicles derived from epithelia appear to fuse dendrites emerging as a potential effector that could repair broken neurons in heterologous systems.

## Materials and Methods

### Strains and transgenic animals

All nematode strains were maintained according to standard protocols [100, 101]. In addition to the wild-type strain N2, the following mutations, transgenes and strains were used: BP601 *aff-1(tm2214)/mIn1[dpy-10(e128) mIs14] II* [55], MF190 *hmIs4[DES-2::GFP, pRF4],* BP328 *eff-1(ok1021) II; hmIs4*, BP450 *hyEx30[myo-2::gfp, DES-2::GFP, KS],* BP431 *eff-1(hy21) II; hmIs4* [33], NC1841 (*wdIs52, F49H12.4::gfp*; *rwIs1*, *pmec-7::*RFP) [34], CHB392 *[hmnEx133(ser-2prom3::kaede)],* kindly provided by Candice Yip and Max Heiman [65]. Germline transformation was performed using standard protocols [102]. The KS bluescript plasmid was used as carrier DNA. Transgenic lines include: BP709 *[hmnIs133 (ser-2prom3::kaede)], BP1014 aff-1/mln1; dzIs53[pF49H12.4::mCherry]Is* was created by crossing *aff-1/ mln1*with *pF49H12.4::mCherry* (kindly provided by Salzburg Y.). BP1015 *aff-1/mln1; hmnIs133[ser-2prom3::kaede]; hyEx66 [KS, pCFJ90 (myo-2::mcherry), pME4(des-2::AFF-1)].* BP1017 *aff-1/mln1; hmnIs133[ser-2prom3::kaede]; hyEx350[KS, pCFJ90 myo-2::mcherry, pTG5 dpy-7p::aff-1],* BP1052 *aff-1/mln1; hmnIs133[ser-2prom3::kaede]; hyEx355[KS, pCFJ90 myo-2::mcherry, pTG4 grd-10p::aff-1],* BP1055 *dzIs53[F49H12.4p::mCherry]; hyEx66[pRF4, AFF-1fosmid::GFP, KS],* BP1056 *dzIs53[F49H12.4p::mCherry]; hyEx68[pRF4, AFF-1 fosmid::GFP, KS].*

### Molecular Biology

We used RF cloning to insert the *grd-10* promoter upstream to the *aff-1* gene [103], and Gateway cloning [104] to clone *aff-1* into a plasmid containing the *dpy-7* promoter fragment (pDest Dpy7 and pDONR™221). Phusion Hot Start II High-Fidelity DNA polymerase (Thermo Scientific, Waltham, MA) was used to facilitate the cloning process.

### Confocal microscopy and live imaging *of C. elegans*

Nematodes were mounted on 3% agar pads mixed with 10 mM NaN_3_ in M9 buffer. For time-lapse analysis, worms were anesthetized with 0.1% tricaine and 0.01% tetramisol in M9 solution [105–107]. Animals were analyzed by Nomarski optics and fluorescence microscopy, using Zeiss LSM 510 META confocal, the Zeiss LSM 700 confocal or Nikon eclipse Ti inverted microscope equipped with Yokogawa CSU-X1 spinning disk (Yokogawa, Tokyo, Japan) and a sCMOS (Andor, Belfast, UK) camera. Z-stacks were taken with PlanApochromat 60x oil NA=1.4 objective using the SDC or 63x NA=1.4 objective using the LSM. When using the sCMOS (Andor) camera z-stacks were taken with ~0.35 µm z-step. When the LSM 510 meta was used, z-step was ~0.8 µm. Image acquisition was done using Andor iQ or Metamorph software (Molecular Devices, Sunnyvale, CA) when using the spinning disk confocal (SDC), and Zen software when using the LSM 510 meta microscope or Zeiss LSM 700. Multidimensional data was reconstructed as projections using the ImageJ and Metamorph softwares. Figures were prepared using ImageJ, Adobe Photoshop CS5 and Adobe Illustrator CS6.

### Quantifying PVD branching phenotypes and statistics

Quantification was done as previously described [33]. Using confocal microscopy, at least five sequential z-series pictures were taken from each worm. Each z-section was analyzed separately. The results from each worm were normalized to a longitudinal length of 100 μm in all relevant experiments. Significant differences between mutants and wild-type were determined by the two-tailed unpaired t-test, the nonparametric Mann-Whitney test or Fischer’s exact test. For each group we observed >20 additional animals that were not recorded by z series on the confocal microscope and that showed similar phenotypes.

### Laser surgery

Microsurgery was done using the LSM 510 META and a tunable multiphoton Chameleon Ultra Ti-Sapphire laser system (Coherent, Santa Clara, CA), that produces 200-fs short pulses with a repetition rate of 113 MHz and 5 nJ energy at a wavelength of 820 nm. 0.5-2 μm^2^ selected rectangular ROIs of GFP-labeled PVD neurons were cut and successful surgery was confirmed by visualizing targets immediately after exposure. We evaluated the ability of severed neurons to reconnect by analyzing z-stack images of GFP-labeled branches. Dendrotomy was performed on the primary longitudinal process, and the morphological changes were followed for 2 to 72 hours after the surgery. Imaging before and after surgery was done as described above, using the 488 nm line of the Argon laser of the LSM microscope or using the spinning disk confocal system. After surgery, animals were recovered to an agar plate and remounted 5-72 hours after surgery. Recovered worms were analyzed for regeneration, fusion between processes, and ectopic sprouting. At least 10 individuals were observed for each experiment. For all worms the primary dendrite was injured anterior to cell body. Animals were imaged and a z-stack was collected immediately after injury to confirm a successful injury.

### Photoactivation using Kaede

In order to verify that dendrites fuse as response to injury we used the photoconvertible protein Kaede driven by a PVD specific promoter *ser-2prom3* [65]. A PVD primary dendrite of *ser-2prom3::Kaede* expressing animals was dendrotomized, animals were recovered for 23 hours and the dendrite reconnection to its stump was assessed by Kaede photoconversion [66]. The green Kaede form in the PVD cell body was irreversibly photoconverted to the red Kaede form using a 405 nm laser with the Mosaic system (Andor) on the Nikon eclipse Ti inverted microscope. Following photoconversion of the Kaede in the cell body we followed spreading to the dendritic branches for 1 and 60 min post photoconversion. Red Kaede form, though diluted while spreading through the dendritic tree, can be observed beyond the reconnection site of injury in the distal part of the primary and higher ordered dendritic branches. When the dendrites fail to reconnect or immediately after dendrotomy the photoconverted Kaede did not cross the site of injury, revealing that spreading of red Kaede is a reliable tool to confirm rejoining of severed dendrites.

## Acknowledgements

We thank A. Fire for vectors, M. Krause for *myo-2::gfp,* the *C. elegans* knockout consortium for the *eff-1(ok1021)* deletion allele, M. Heiman and C. Yip for CHB392, C. Smith and D. Miller for NC1841. We also thank A. Sapir, B. Gildor, and J. Grimm for critical comments on earlier versions of the manuscript, and E. Oren for MATLAB advice. Some strains were provided by the CGC, which is funded by NIH Office of Research Infrastructure Programs (P40 OD010440).

## Author contributions

Conceived and designed the experiments: MOS TG BP. Performed the experiments: MOS TG VK. Analyzed the data: MOS TG BP. Wrote the paper with input from all authors: MOS BP.

## Supporting information captions

**S1 Fig. Dendritic amputation induces hyper-branching, loss of self-avoidance and giant overlapping menorahs**

**(A)** Wild-type animal before dendrotomy.

**(B)** Two laser-induced injuries of primary branches (red arrowheads, injury sites).

**(C)** Confocal projection and a schematic tracing of the same animal 24 hr post-surgery. Asterisks mark new sprouting from proximal and distal branches. Pink brackets, giant menorahs. Blue arrowheads, primary branch fusion.

**(D)** A different wild-type animal injured (red arrowhead) at the early L4 stage, recovered and analyzed 24 **(E)** and 46 hr **(F)** post-surgery. Black arrow, retrograde branch. The degenerating branches are represented as dotted lines in the schematic tracing. Scale bars represent 5 µm **(D)** and 10 µm **(A-C, E-F)**.

**S2 Fig. Menorah fusion and 2ry stem pruning are part of the regeneration process of the PVDs following dendrotomy**

**(A)** Dendrotomy of L4 wild-type animal. Red arrowheads point to sites of laser surgery.

**(B)** Analysis 48 hr post-surgery. Giant menorah can be seen (pink bracket); arrows (and dotted lines in the schematic tracing) point to secondary branches undergoing trimming. Blue arrowheads mark primary branch fusion at the sites of dendrotomy. Scale bars represent 10 µm.

**S3 Fig. PVD dendrite reconnection confirmed by Kaede photoconversion**

A PVD primary dendrite of *ser-2prom3::Kaede* expressing animals was dendrotomized, the animal was recovered for 23 hr and the dendrite reconnection to its stump was assessed by Kaede photoconversion. The green Kaede form in the PVD cell body was irreversibly photo-converted to the red Kaede form using a 405nm laser with the Mosaic system and its spreading throughout the dendritic branches was followed 1 and 60 min post-photoconversion. Panels left to right are confocal reconstructions of a wild-type dendrotomized animal in the 488 green channel, 561 red channel, two channels merged view and a schematic representation of the merged view.

**(A)** Confocal reconstructions of the animal before dendrotomy, **(B)** immediately post dendrotomy, **(C)** 23 hr post-dendrotomy, **(D)** 1 min post Kaede photoconversion and **(E)** 60 min post Kaede photoconversion. Green Kaede (magenta in merged and schematic representation). Red Kaede form (cyan in merged and schematic representations), though diluted when spreading through the dendritic tree, can be observed beyond the reconnected site of injury in the distal part of the primary and higher ordered dendritic branches. Thus, Kaede photoconversion and diffusion beyond the injury site demonstrate fusion between the severed primary dendrite. In animals where reconnection failed photoconverted Kaede did not spread beyond the injury site (data not shown).

c, PVD cell body; red arrowhead, site of injury; blue arrowhead, site of primary branch fusion. In the merged and schematic columns: magenta, green Kaede; cyan, photoconverted red Kaede.

**S4 Fig. Ectopic expression of AFF-1 does not improve PVD regeneration in heterozygous animals**

**(A)** Regeneration by menorah-menorah fusion following PVD nanosurgery in *C. elegans aff-1(tm2214)/mln1; dpy-7p::aff-1* animals. PVD is shown before and after nanosurgery (T=0, 3, and 4 hr). PVD reconnection occurred through fused menorahs (pink bracket) and primary fusion did not occur (red arrowhead).

**(B)** Dendrite regeneration following PVD primary branch nanosurgery in *C. elegans aff-1(tm2214)/mln1; dpy-7p::aff-1* animals. PVD is shown before and after nanosurgery (T=0 and 2 hr). PVD reconnection occurred through primary fusion (blue arrowhead).

**(C)** Animals showing different PVD post injury consequences displayed in color coded pie graphs as Magenta-Menorah-Menorah fusion, Blue-primary-primary fusion, Magenta and Blue-Menorah-Menorah fusion and primary-primary fusion, Black-degeneration. Wild type n=22; *aff-1(tm2214)* n=32; *aff-1/mln1; des-2p::AFF-1* n=10, *aff-1(tm2214) /mln1; grd-10p::aff-1* n=11, *aff-1(tm2214)/mln1; dpy-7p::aff*-1 n=14. *P<0.05. P values for wild type and heterozygous genotypes are not significant (NS). Statistics was calculated using the Freeman-Halton extension of the Fisher exact probability test for a two-rows by four-columns contingency table, <u>http://vassarstats.net/fisher2x4.html</u>.

Dendrotomy site, red arrowhead; fused Menorah, pink bracket; primary fusion, blue arrowhead. Scale bars represent 20 µm.

**S5 Fig. Model of AFF-1-mediated repair via extracellular vesicle-cell fusion** PVD (red) is in close proximity to the epithelial seam cells (blue). AFF-1 (black pins) is expressed in seam cells and additional tissues, but not in the PVD. Upon injury, AFF-1-containing extracellular vesicles (EVs) are highly released from the seam cells. Some of these EVs reach the PVD and promote fusion of severed dendrites. EFF-1 (green pins) is expressed in the PVD but it does not act to fuse severed dendrites on its own. Instead it may collaborate with AFF-1-EVs. We propose that menorah-menorah fusion is mediated by AFF-1-EVs that merge with the structurally compatible EFF-1 expressed in the PVD. EFF-1-coated pseudotyped viruses can fuse with cells expressing AFF-1 on their surface and vice versa [68].

## Supplemental movies captions

**Movie 1. PVD dendrites touch and fuse following dendrotomy**

Time lapse recording of an L4 wild-type animal just after injury. The z series images were recorded every 6 min, marker is *F49H12.4::GFP.* Arrows mark areas of fusing menorah tertiary branches. C marks PVD cell body.

**Movie 2. A pseudo-colored presentation of depth information for Movie 1**

Scale bar for the position in the z axis is shown at the bottom. The movie was obtained using the Zeiss LSM image browser DepthCod function.

**Movie 3. Degeneration of the distal fragment following dendrotomy**

Time lapse recording of a wild-type L4 animal after a two-photon injury. Posterior PVCR and PVCL are also marked with the *DES-2::GFP* marker and can be seen at the upper right corner. See **Fig 1G**.

**Movie 4. Fusion and pruning during the PVD regeneration process**

Intensity-values view of a time-lapse recording of an early L4 animal. Marker is *F49H12.4::GFP.* Menorahs from proximal and distal ends meet and fuse, bypassing the break induced by the two-photon injury. At the injury site growth and pruning of dendrites can be seen. Intensity scale bar is in **Fig 2D**.

**Movie 5. AFF-1::GFP expression in PVD intact animals**

Time lapse recording of *AFF-1::GFP* (cyan) expression inside and outside the seam cells (sc), and in the vulva (V) of an intact L4 animal shown in a two channels merged image. The z series images were recorded every 10 sec, PVD marker is *F49H12.4p::mCherry* (magenta). Arrows mark AFF-1 vesicles. See **Fig 4B**. The trajectory of vesicle 1 is shown in **Fig 4E**. Gut marks autofluorescent gut granules, easily distinguishable from AFF-1 vesicles in size, shape, and appearance in both channels.

**Movie 6. AFF-1::GFP expression in PVD intact animals (zoom in of movie 5)**

A magnification of a single *AFF-1::GFP* vesicle near the seam cells, (“vesicle 1”).

**Movie 7. AFF-1::GFP expression in PVD primary branch of injured animals**

Time lapse recording of *AFF-1::GFP* expression (cyan) inside and outside the seam cells (sc), in the anchor cell (AC) and in the vulva (V) of a reconnected PVD L4 animal after a two-photon injury anterior to PVD cell body. T=0 is 75 min after injury. The z series images were recorded every 20 sec, PVD marker is *F49H12.4p::mCherry* (magenta). Arrows point to vesicles containing AFF-1::GFP.

**Movie 8. Magnification of Movie 7**

Magnification of an area with multiple vesicles, two are labeled (2,3). See **Fig 4 D-E.**

**Movie 9. AFF-1::GFP expression in PVD intact animal**

Time lapse recording of *AFF-1::GFP* (cyan) expression inside and outside the seam cells (SC) of an intact L4 animal shown in a two channels merged image. The z series images were recorded every 10 sec, PVD marker is *F49H12.4p::mCherry* (magenta). Arrows mark AFF-1 vesicles.

**Movies 10 and 11. AFF-1::GFP expression in a degraded PVD primary branch of injured animals**

Time lapse recording of *AFF-1::GFP* expression (cyan) inside and outside the seam cells (sc) of a reconnected PVD L4 animal after a two-photon injury anterior to PVD cell body, red arrowhead is the site of injury. T=0 is 75 min (movie10) and 150 min (movie 11) after injury. The z series images were recorded every 2 sec, PVD marker is *F49H12.4p::mCherry* (magenta). Arrows point to vesicles containing AFF-1::GFP.

